# Social groups with diverse personalities mitigate physiological stress in a songbird

**DOI:** 10.1101/2020.04.15.043471

**Authors:** Csongor I. Vágási, Attila Fülöp, Gergely Osváth, Péter L. Pap, Janka Pénzes, Zoltán Benkő, Ádám Z. Lendvai, Zoltán Barta

## Abstract

Social groups often consist of diverse phenotypes, including personality types, and this diversity is known to affect the functioning of the group as a whole. Social selection theory proposes that group composition (i.e. social environment) also influences the performance of individual group members. However, the effect of group behavioural composition on group members remains largely unexplored, and it is still contentious whether individuals benefit more in a social environment with homogeneous or diverse behavioural composition. We experimentally formed groups of house sparrows *Passer domesticus* with high and low diversity of personality (exploratory behaviour), and found that their physiological state (body condition, physiological stress and oxidative damage) improved with increasing group-level diversity of personality. These findings demonstrate that group personality composition affects the condition of group members and individuals benefit from social heterosis (i.e. associating with a diverse set of behavioural types). This aspect of the social life can play a key role in affiliation rules of social animals and might explain the evolutionary coexistence of different personalities in nature.

## 1. Introduction

Social groups usually consist of a mixture of members with diverse phenotypes [1]. Variation within groups can occur in morphology (e.g. size), behavioural traits (e.g. reactive and proactive behavioural types [2], or social roles (e.g. leaders/followers in human *Homo sapiens* teams [3], or producers/scroungers in tree sparrow *Passer montanus* flocks [4]). Group composition has implications for emergent group-level processes such as decision-making, which ultimately drive group functioning (reviewed in [1,5]). Ethnic diversity can, for instance, have a positive effect on research teams’ scientific performance [6].

Personality, the consistent among-individual differences in behavioural phenotype [7], has strong relevance for social life [8]. Social groups can largely differ in their personality composition some being more homogenous, while others more heterogeneous [1]. Groups’ personality composition can have effects at both the level of group as a whole (‘upstream effects’) and the level of individual group members (‘downstream effects’). The group-level consequences of groups’ personality composition have mostly been assessed in human teams [9,10], where personality composition may affect team performance, albeit in an inconsistent manner [11–13]. In non-human organisms, group personality composition influences within-group social network structure, collective behaviour, and group performance [5,14–17]. Although individual-level aspects of sociality, e.g. rank in the dominance hierarchy, have been shown to influence the state of individual members (reviewed by [18,19]), the downstream effects of group behavioural composition on individual state is surprisingly little scrutinised [1]. This happens despite that social selection theory postulates that the fitness of an individual is contingent upon the phenotype of those with whom it affiliates (i.e. social environment; [15,20]).

Health state can have a strong influence on the individual’s social behaviour [21] and, at the same time, is moulded by changes of individual’s social environment [19]. Therefore, it is reasonable to assume that physiological condition (e.g. body condition, stress physiology, oxidative damage and immune capacity) of group members is also influenced downstream by the personality composition of their group. Correlative studies on human work teams found that age and gender composition can be associated with subjective self-reported health impairment [22], but human studies with experimental manipulation of group composition and actual health measurements are still lacking. Earlier animal studies mostly assessed how individuals’ position within the social structure (e.g. rank in dominance hierarchy, an individual-level social attribute) affected their health or physiology (reviewed by [18,19]). Experimental studies on animals that tested directly whether group behavioural composition (a group-level social attribute) affects the stress level and condition of group members are very scarce and each involve livestock species [23,24]. No experimental study addressed this question in wild animals.

How could group composition affect the state of group members? Several non-exclusive mechanisms can play a role. Diverse groups provide more opportunities for specialization [1,25] and are more likely to host keystone individuals, which are influential individuals with disproportionately large effect on other group members and/or overall group functioning [26]. Both role specialization and keystone individuals can lead to superior group-level performance (upstream effect). Indeed, great tit *Parus major* affiliations consisting of diverse personalities show the most effective coordinated action when exploring a habitat patch [5]. Similarly, a mixture of shy and bold guppies *Poecilia reticulata* can be advantageous in reducing the trade-off between exploring novel foraging tasks and antipredator vigilance [16]. The minority of keystone individuals can also substantially affect group-level behaviour and performance [14].

Complementarity of different personalities might also be advantageous in groups of diverse personalities. Stickleback *Gasterosteus aculeatus* shoals solve better a two-stage food acquisition problem when the shoal contains fish that have experience with stage one and fish that have experience with stage two (termed “experience-pooling”; [27]). Finally, groups with diverse behavioural composition might experience less aggression in pigs *Sus scrofa domesticus* [23]. These group-level advantages of diversely composed groups can bring about higher individual performance (downstream effect) in terms of either fitness or condition [17] by reducing stress exposure and ultimately leaving group members in better physiological condition. Here we asked whether group-level diversity of personalities might influence the physiological state of individual group members.

We conducted an experimental study to explore whether manipulated personality composition of groups *per se* or in interaction with individual personality type affects the physiological condition of group members. Given that diverse groups might have advantages for group functioning (see above) and potentially there is less aggression in behaviourally diverse groups [23], we predicted that individuals in groups with more diverse personality composition would improve their physiological condition as compared with individuals in groups with more homogeneous personality composition. In this case, improved physiological condition of group members is a legacy of being part of a diversely composed and better functioning group and each group member shares this legacy. We do not know whether this potential benefit is indeed uniform for each member. Alternatively, some members might harvest more the benefits of better group-level functioning to the detriment of other group members. We assessed this prediction by testing whether personality diversity of the groups interact with individual personality type to influence physiological condition. A significant interaction might suggest that individuals that either match or mismatch the group’s personality composition benefit more than other group members do.

## 2. Material and methods

### (a) Study species

The house sparrow *Passer domesticus* is arguably one of the most popular model organisms in animal ecology and evolution [28]. It is an ideal candidate to study sociality, since it exhibits a wide spectrum of social behaviour including colonial breeding, social foraging or communal roosting [29]. At our study site, house sparrows are year-round residents, breed in cavities of stall buildings at the cattle farm and forage in flocks of various sizes, especially outside the breeding season. Flock sizes vary from small to medium, containing from a few birds up to some dozen individuals similarly to other populations (see [29]).

### (b) Study protocol

The study is based on a large sample of 240 house sparrows. The same study protocol was used in six study replicates. We captured 40 sparrows (1:1 sex ratio) per each study replicate (Electronic Supplementary Material, ESM table S1). These 40 birds were divided into four treatment groups (see below) consisting of 10 birds each, which yielded 24 social groups for the entire study (four treatment groups per study replicate × six study replicates). There was no significant difference in sex ratio between the treatment groups in any of the study replicates (χ^2^ test, all *p* > 0.362).

The study timeline in each study replicate was as follows. Upon capture (day 0), birds were marked with an aluminium ring, and their sex and body mass was recorded. Then they were housed in indoor aviaries for 18 days at the campus of Babe□-Bolyai University, Cluj-Napoca, Romania. On days 5–7, we recorded exploratory behaviour as a well-established and ecologically relevant axis of personality [30] following the novel environment test of Dingemanse et al. [31]. At day 9, we measured the body mass and tarsus length of the birds, and took a pre-treatment blood sample. Then the birds were allocated according to an *a priori* defined, stratification based protocol (for details, see ESM) into one of four social treatment groups of 10 birds each: ‘random’ (random subsample of birds of a given replicate), ‘low exploratory’ (only birds with low exploratory scores), ‘high exploratory’ (only birds with high scores), and ‘variable’ (equal mixture of birds with either low or high scores) (see details of Social treatment methods in ESM). Social treatment protocol successfully created differences in the mean and variance of exploration score (personality) among groups as recommended by Farine et al. [1] (ESM figure S1). The mean exploratory score was the smallest in the low exploratory group, intermediate in the random and variable groups, and the largest in the high exploratory group (Linear mixed-effects model with study replicate as random factor, treatment group effect: χ^2^ = 632.92, df = 3, *p* < 0.001; figure S1a). The variance of exploratory score was the lowest in the low and high exploratory groups, intermediate in the random group and the highest in the variable groups (treatment group effect: χ^2^ = 401.78, df = 3, *p* < 0.001; low vs. high exploratory: χ^2^ = 1.93, df = 1, *p* = 0.165; random vs. variable: χ^2^ = 58.58, df = 1, *p* < 0.001; figure S1b). The significant difference in variance of exploratory score between the random and the variable groups is due to the former being an even sample of the entire range of exploratory behaviour, while the latter being a mixture of low exploratory and high exploratory individuals in equal proportion (see details of Social treatment methods and figure S2 in ESM). Therefore, as an additional characterisation of the birds’ social environment, we calculated *a posteriori* the Shannon diversity index of exploratory behaviour for each social group of 10 sparrows by dividing exploration values into 10 ordered categories of roughly equal sizes. As expected, the Shannon diversity index was the lowest in the low and high exploratory groups, intermediate in the variable group, and the highest in the random group (treatment group effect: χ^2^ = 68.118, df = 3, *p* < 0.001; low vs. high exploratory: χ^2^ = 1.30, df = 1, *p* = 0.254; random vs. variable: χ^2^ = 11.73, df = 1, *p* < 0.001; figure S1c).

The social treatment period lasted nine days until day 18, when we measured again the body mass and took a second blood sample to measure the post-treatment physiological condition. Physiological state was characterised by measuring body condition (scaled body mass index, SMI), heterophil-to-lymphocyte (H/L) ratio, oxidative damage to lipids (malondialdehyde concentration, MDA), and innate immune capacity (natural antibodies – agglutination score; complement system – lysis score) both during the pre- and post-treatment sampling events. These traits were chosen for the following reasons. Changes in body condition usually take more time than changes in other physiological traits. Among the factors that affect body condition, exposure to stress stimuli is known to reduce body condition [32]. Ultimately, impaired body condition can have widespread consequences for the organism. H/L ratio has been shown to correlate with the glucocorticoid stress response governed by hypothalamic-pituitary-adrenal axis [33,34], but is less sensitive to handling stress as compared with plasma levels of corticosterone (the main glucocorticoid in birds). H/L ratio therefore can indicate the degree of physiological stress. MDA is a versatile marker of oxidative stress for two reasons. MDA indicates the damage to vital cell membrane lipids, which has substantial adverse consequences from the cell level to the organism level. Besides being a direct measure of oxidative damage, MDA is a pro-oxidant itself with a long half-life and hence can reach far from its site of origin damaging other vital macromolecules [35]. Agglutination and lysis capacity of the plasma describes the activity of the humoral innate immune system and hence is an indicator of the first line of defence of vertebrate hosts against invading microorganisms [36]. Higher scores of agglutination and lysis indicate higher innate immune responsiveness. There was no significant difference among social treatment groups in the pre-treatment values of the five physiological response variables (all *p* > 0.188; see detailed statistics in ESM Results). The different physiological variables are weakly correlated both in the pre- and post-treatment samples except for the positive association between agglutination and lysis scores (table S2). We provide a more detailed description of study timeline, captivity conditions, measurement of exploratory behaviour, assignment to social treatment groups, blood sampling and physiological measurement methods in the ESM.

### (c) Statistical procedures

The statistical analyses were carried out in the R statistical environment (ver. 4.03; [37]). Since we used a repeated-measures approach to analyse our data, each physiological variable contained the values of both sampling events (i.e. pre-treatment and post-treatment values). Each physiological trait was analysed in a separate statistical model. Heterophil-to-lymphocyte ratio was arcsine square root transformed, malondialdehyde level was log-transformed and exploration score was log(x+1)-transformed to reduce the bias in their distributions, while haemagglutination and haemolysis scores were converted into binary variables (0 for absence and 1 for presence of agglutination or lysis) because they were highly zero-inflated. To improve model convergence, the continuous dependent variables (i.e. body condition, heterophil-to-lymphocyte ratio and malondialdehyde concentration) and the continuous predictor variables (exploration score and Shannon diversity) were Z-transformed to have zero mean and unit standard deviation [38].

In the first set of models, we assessed the effect of social treatment and the effect of sampling event × social treatment interaction on the individual physiological responses of sparrows. The explanatory variables were the same in all models as follows: sex (male or female), social treatment (random, variable, high-exploratory and low-exploratory) and sampling event (pre-treatment and post-treatment) were set as fixed factors, and individual exploratory score as a continuous predictor. In addition, all second-order interactions between the four explanatory variables were also tested. Note that, in this setup, a significant interaction with sampling event indicates that the rate of change in the response variable is influenced by the other explanatory variable in the interaction. Study replicate, treatment group ID nested within study replicate, and individual ID nested within study replicate and treatment group ID were entered as random factors. We used linear mixed-effects models with normal error distribution (LMMs; ‘lmer’ function of the R package ‘lme4’; [39]) for body condition, heterophil-to-lymphocyte ratio and malondialdehyde, while we used generalized linear mixed-effects models with binomial error distribution (GLMMs; ‘glmer’ function of the R package ‘lme4’) for agglutination and lysis scores. The assumption of homogeneity of residual variances among treatment groups were met for each response variable of the LMM models (Levene test, all *p* > 0.195). We assessed the fulfilment of model assumptions by graphical diagnosis; all assumptions were met for each model. Each model was simplified to obtain minimal adequate models (MAMs) containing only significant main effects or their interactions by sequentially dropping predictors with non-significant (*p* > 0.050) effects using the ‘drop1’ R function. The sampling event × social treatment interaction and its main effects were kept in the model even if they were non-significant because our main interest is related to the sampling event × social treatment interaction.

In the second set of models, we assessed the effect of Shannon diversity of group personality and the effect of sampling event × Shannon diversity interaction on the individual physiological responses of sparrows. For this, we used a similar approach as in the first model set with the only difference that we entered groups’ Shannon diversity of exploration as a continuous predictor in the models instead of the fixed effect of social treatment.

The reported significance levels were calculated using type II Wald Chi-square tests using the ‘Anova’ function of the R package ‘car’ [40]. The post-hoc comparisons of individual changes in physiological variables between the two sampling events as a function of different treatment groups (first set of models) or as a function of groups’ Shannon diversity of exploration (second set of models) were conducted using the R package ‘emmeans’ (functions ‘emmeans’ and ‘emtrends’, respectively; [41]). Tables 1 and 2 present the type II ANOVA results of MAMs for the first and second set of models, respectively. ESM tables S2 and S3 present the parameter estimates of both the full models and the MAMs for the first and second set of models, respectively. All data and analyses code are deposited in Dryad Digital Repository [42].

**Table 1.** Minimal adequate models containing predictors of individual responses in physiological state of house sparrows to social treatment. Statistically significant effects (*p* ≤ 0.050) are marked in bold, while marginally significant effects (0.050 < *p* ≤ 0.100) in italic. SMI – Scaled Mass Index (body condition), H/L ratio – heterophil-to-lymphocyte ratio (physiological stress response), MDA – malondialdehyde (oxidative damage to lipids).

| response | fixed effects | $\chi^2$ | df | $p$ |
| --- | --- | --- | --- | --- |
| SMI | sampling event | 0.00 | 1 | 1.000 |
|  | treatment | 3.70 | 3 | 0.296 |
|  | **sex** | **13.66** | **1** | **< 0.001** |
|  | **sampling event × treatment** | **14.32** | **3** | **0.003** |
| H/L ratio | sampling event | 0.00 | 1 | 1.000 |
|  | treatment | 1.38 | 3 | 0.709 |
|  | sex | 0.02 | 1 | 0.879 |
|  | *sampling event × treatment* | *7.56* | *3* | *0.056* |
|  | **sampling event × sex** | **3.85** | **1** | **0.050** |
| MDA | sampling event | 0.00 | 1 | 0.979 |
|  | treatment | 1.04 | 3 | 0.792 |
|  | **sampling event × treatment** | **13.83** | **3** | **0.003** |
| agglutination | **sampling event** | **12.54** | **1** | **< 0.001** |
|  | treatment | 1.51 | 3 | 0.680 |
| | sampling event × treatment | 0.81 | 3 | 0.848 |
| lysis | **sampling event** | **16.17** | **1** | **< 0.001** |
|  | *treatment* | *7.09* | *3* | *0.069* |
| | sampling event × treatment | 0.38 | 3 | 0.945 |

**Table 2.** Minimal adequate models containing predictors of individual responses in physiological state of house sparrows to social treatment in relation with the groups’ personality diversity. Statistically significant effects (*p* < 0.050) are marked in bold. SMI – Scaled Mass Index (body condition), H/L ratio – heterophil-to-lymphocyte ratio (physiological stress response), MDA – malondialdehyde (oxidative damage to lipids).

| response | fixed effects | $\chi^2$ | df | $p$ |
| --- | --- | --- | --- | --- |
| SMI | sampling event | 0.00 | 1 | 1.000 |
|  | **sex** | **12.68** | **1** | **< 0.001** |
|  | Shannon diversity | 1.57 | 1 | 0.210 |
|  | **sampling event × Shannon diversity** | **14.44** | **1** | **< 0.001** |
| H/L ratio | sampling event | 0.00 | 1 | 1.000 |
|  | Shannon diversity | 0.00 | 1 | 0.977 |
| | sampling event × Shannon diversity | 0.62 | 1 | 0.431 |
| MDA | sampling event | 0.00 | 1 | 0.983 |
|  | Shannon diversity | 1.37 | 1 | 0.241 |
|  | **sampling event × Shannon diversity** | **6.10** | **1** | **0.014** |
| agglutination | **sampling event** | **12.36** | **1** | **< 0.001** |
|  | Shannon diversity | 0.18 | 1 | 0.675 |
| | sampling event × Shannon diversity | 1.49 | 1 | 0.222 |
| lysis | **sampling event** | **16.10** | **1** | **< 0.001** |
|  | Shannon diversity | 0.02 | 1 | 0.882 |
| | sampling event × Shannon diversity | 0.10 | 1 | 0.756 |

## 3. Results

### (a) Social treatment effects

In the first model set, we assessed the effects of social treatment on the individual’s physiological responses. The sampling event × social treatment interaction was significant for body condition and oxidative damage, and marginally significant for heterophil-to-lymphocyte ratio, while it was non-significant for agglutination and lysis (table 1, table S3, figure 1; see also the individual reaction norms of physiological traits to social treatment on figure S3). The main effect of social treatment was non-significant for each response variable, except for a marginal effect in case of the complement system of the innate immunity (i.e. lysis score; table 1, table S3). Below we present the results related to the physiological variables for which the sampling event × social treatment interaction was significant.

**Figure 1.**
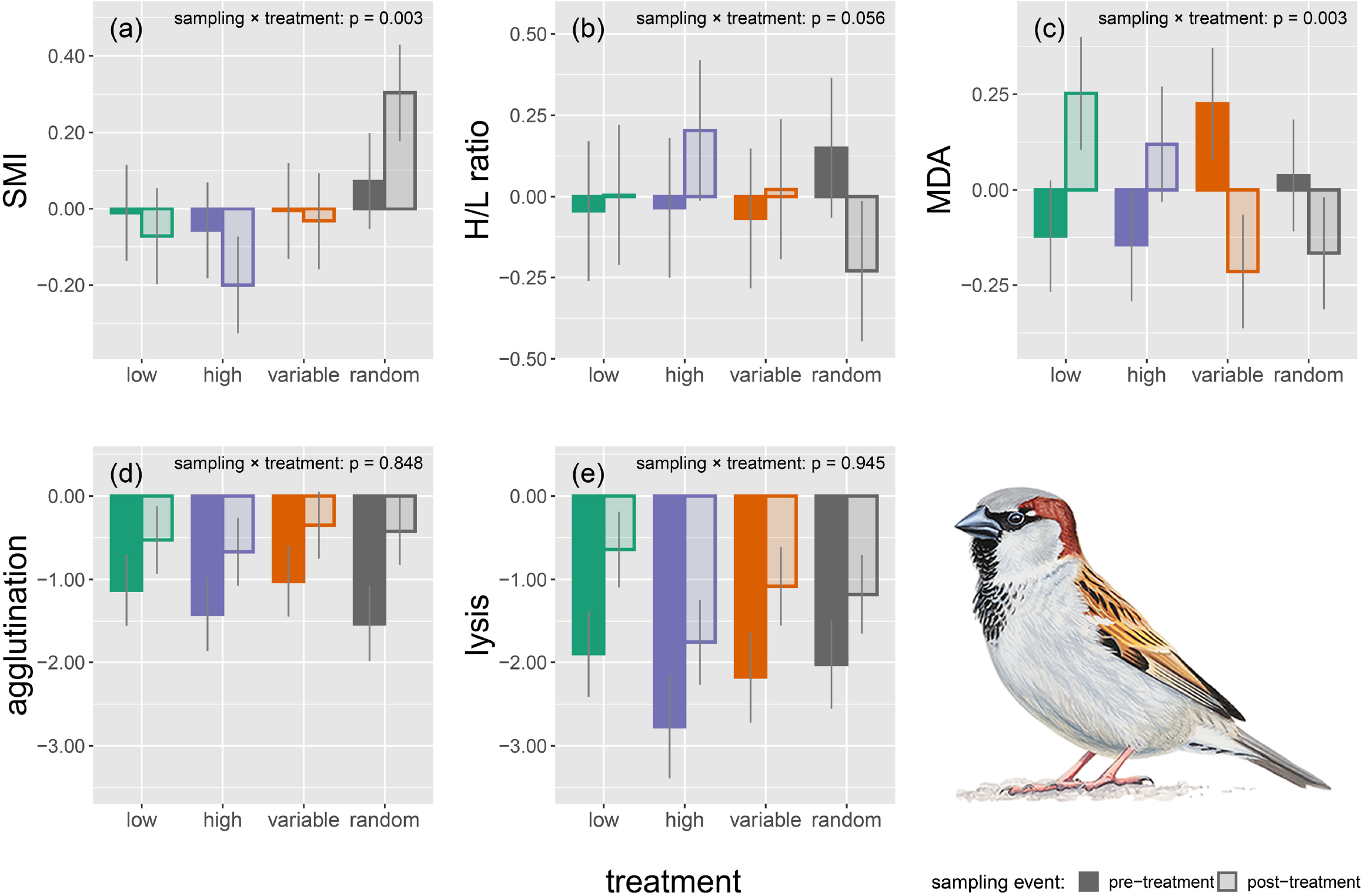
Physiological responses to social treatment: (a) body condition (SMI), (b) heterophil-to-lymphocyte ratio (H/L ratio, an indicator of physiological stress response), (c) oxidative damage to lipids (i.e. malondialdehyde, MDA), and (d–e) constitutive immune capacity, as expressed through agglutination score (d) and lysis score (e). Model-predicted mean ± s.e. are shown for each treatment group both at pre-treatment (dark bars) and post-treatment (light bars) sampling events. Male house sparrow painting credit: Márton Zsoldos.

The improvement in body condition during the treatment period was significantly higher in the random group as compared with the other three groups (see sampling event × social treatment interaction in table S3a). Post-hoc tests revealed that body condition improved significantly in the random group between the two sampling events (*β* = –0.232, s.e. = 0.074, df = 236, *t* = 3.123, *p* = 0.008), while body condition was not affected by the experimental treatment in the other three treatment groups (all *p* > 0.198). Body condition increased only in the random group and decreased in the other three groups (figure 1a).

The rate of increase in heterophil-to-lymphocyte ratio was significantly higher in the high-exploratory group, and marginally higher in the low-exploratory and variable groups as compared with the random group (see sampling event × social treatment interaction in table S3b). Post-hoc tests showed a weak decrease in heterophil-to-lymphocyte ratio in the random group between the two sampling events (*β* = 0.378, s.e. = 0.166, df = 235, *t* = 2.270, *p* = 0.093), while it remained unchanged in the other three groups (all *p* > 0.484). Heterophil-to-lymphocyte ratio (an indicator of physiological stress) decreased only in the random group, while increased in the other three groups (figure 1b).

The increase in oxidative damage during the treatment period was significantly higher in the low-exploratory group and marginally higher in the high-exploratory group as compared with the random group, while the variable and random groups did not differ (see sampling event × social treatment interaction in table S3c). The post-hoc tests revealed that individuals from the variable group showed a marginally non-significant decrease in oxidative damage levels between the two sampling events (*β* = 0.439, s.e. = 0.179, df = 233, *t* = 2.452, *p* = 0.058), whereas birds from the other three treatment groups were not affected by the experimental manipulation (all *p* > 0.140). Malondialdehyde levels (i.e. oxidative damage to lipids) increased in high- and low-exploratory groups, but decreased in the variable and random groups (figure 1c).

Individuals’ personality was not associated with the individual responses in physiological condition by itself or in interaction with social treatment, hence was dropped from the models (table 1, table S3). Constitutive innate immunity improved during treatment as both agglutination and lysis scores increased significantly between the two sampling events, and males had higher body condition than females (table 1, table S3).

### (b) Shannon diversity of group’s personality

To assess the role of personality diversity of experimental social groups, we tested whether the calculated Shannon diversity index predicted the individual’s physiological responses to social treatment. The sampling event × Shannon diversity interaction was highly significant for body condition and oxidative damage (figure 2), while it was non-significant for heterophil-to-lymphocyte ratio and the activity of constitutive innate immunity as measured by agglutination and lysis (table 2, table S4). The main effect of Shannon diversity was non-significant for each response variable (table 2, table S4). Regarding the significant interactions, we found that body condition increased (*β* = 0.146, s.e. = 0.067, df = 24.9, *t* = 2.195, *p* = 0.038), while oxidative damage to lipids (i.e. malondialdehyde) decreased (*β* = –0.170, s.e. = 0.067, df = 66.0, *t* = 2.532, *p* = 0.014) with increasing personality diversity in the post-treatment sample (i.e. after exposure to social treatment), while these two response variables were unrelated to Shannon diversity in the pre-treatment sample (i.e. prior to social treatment; all *p* > 0.413; table 2, table S4, figure 2). The other three response variables (i.e. heterophil-to-lymphocyte ratio, and agglutination and lysis) were unrelated to Shannon diversity in either the pre-treatment or the post-treatment samples (all *p* > 0.231 and all *p* > 0.578, respectively).

**Figure 2.**
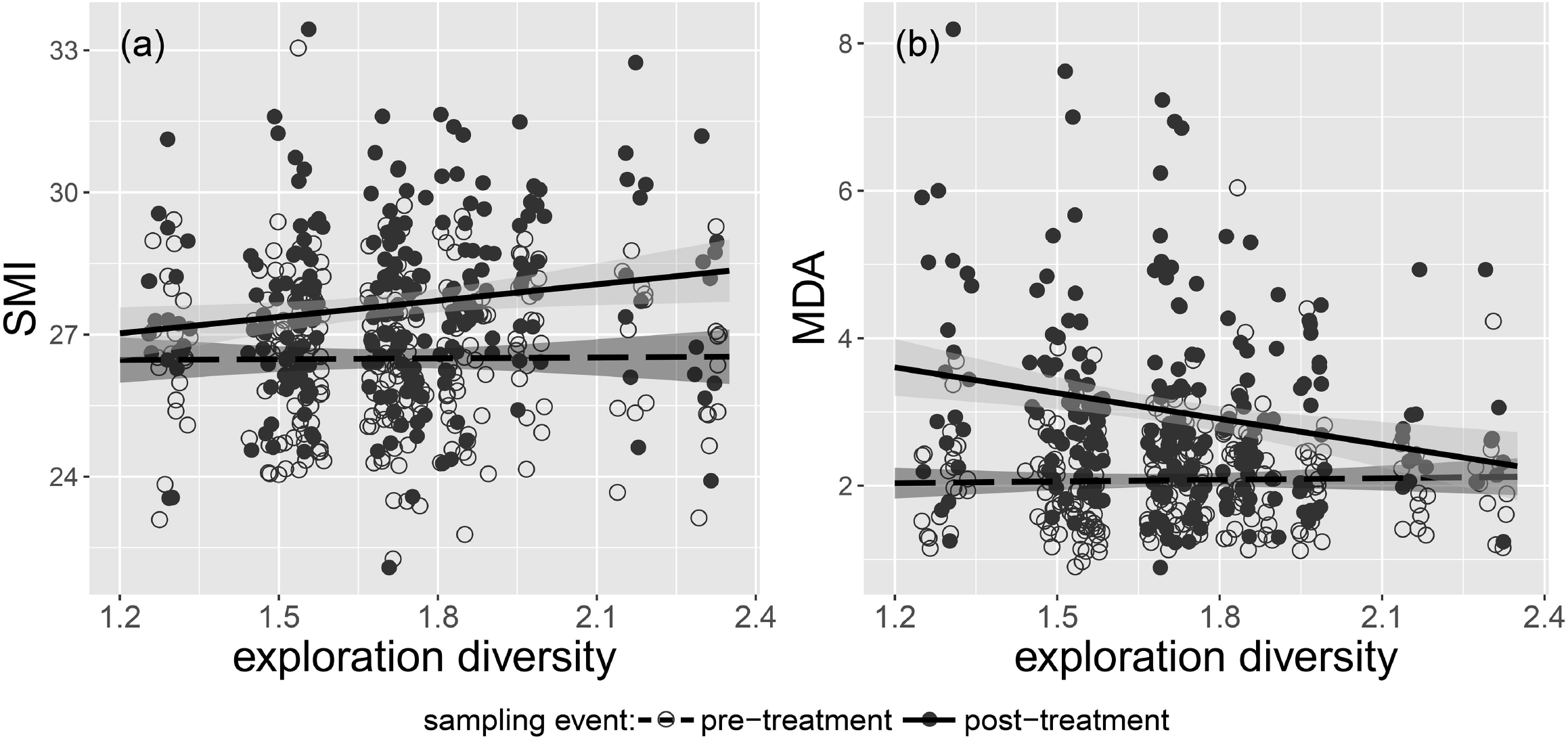
Physiological condition improved in social groups with diverse personalities. Relationships between the Shannon index of group-level personality diversity of house sparrows and individual responses to social treatment in terms of (a) body condition (SMI) and (b) oxidative damage to lipids (MDA). Open symbols and dashed lines denote the pre-treatment sampling, while the filled symbols and continuous lines denote the post-treatment sampling. Regression lines are model-predicted slopes with 95% confidence intervals (shaded areas).

## 4. Discussion

In the realm of human psychology, there is a long-standing debate on whether uniformly or diversely composed teams perform better [11,12], but almost nothing has been known about whether group behavioural composition affects the physiological condition of group members either in humans or other animals. For instance, pigs housed in groups of uniform behavioural composition or in groups with a mixture of behavioural types did not differ in their glucocorticoid stress response or weight gain [23,24]. In contrast, our results are consonant with the old Latin proverb “Varietas delectat” [43], as here we showed that condition significantly improved, heterophil-to-lymphocyte ratio (an indicator of physiological stress) was marginally reduced and oxidative damage level was significantly reduced for house sparrows living in a social environment with diverse personalities. Therefore, living in social groups with diverse composition can provide at least short-term benefits in terms of reduced physiological stress and superior condition. The finding of no significant interactions between social treatment and individual’s exploration score suggests that all individuals in diverse groups similarly enjoy these benefits.

The two heterogeneously composed social treatment groups of our study (i.e. ‘variable’ and ‘random’ groups) had higher diversity of personalities than the two homogeneously composed ones (i.e. ‘low-exploratory’ and ‘high-exploratory’ groups). Nevertheless, the effect of treatment on body condition and heterophil-to-lymphocyte ratio differed between the variable and random groups, as only birds of the random group fared well. This suggests that there is a personality diversity threshold, which has to be exceeded to reap physiological benefits of group composition. Indeed, the heterogeneous variable group only marginally differed from the homogeneous high exploratory group, while the random group stood out among all groups in terms of personality diversity (figure S1c). Therefore, an equal mixture of low and high exploratory birds (i.e. personality extremes) in the variable group is not necessarily sufficient, but a more diverse mixture of the entire range of personalities, like in the random group, is required for some of the individual-level physiological benefits to emerge. A possible explanation for such a threshold effect could be that there is more room for role specialization, keystone individuals and complementarity of different personalities in diversely composed groups [1,25,26], which lead to superior group-level performance (upstream effect) [5,14,16,27] and ultimately precipitate superior individual-level performance (downstream effect) [17]. This hypothesis can be tested by studying social groups that differ in personality diversity succeed in solving novel tasks (e.g. in a foraging context).

It is important to note that we found a positive effect of social diversity on health in a set-up where constant amount of food was available. This suggests that the benefits of social diversity might arise because of the type and intensity of aggressive interactions between members rather than being the consequence of more obvious benefits like improved habitat exploration, defence against predators and decision-making. Therefore, preference for bonding with dissimilar individuals (i.e. heterophily), and the resulting better health state of individuals in diverse groups might reinforce the improved group-level outcomes creating a positive feedback loop [44] between group-level and individual-level performances. Indeed, affiliations of great tits and simulated social groups consisting of diverse personalities show the most effective coordinated action when exploring a habitat patch (group-level performance) [5], while social groups composed of diverse personalities also improve physiological condition (individual-level performance; present study). This simultaneous group- and individual-level benefits of diversely composed groups can drive the evolutionary maintenance of heterophily [45].

Consistency of personality traits places a constraint on individuals because one is either more reactive (shy, neophobic, less exploratory, less aggressive) or more proactive (bold, novelty seeking, more exploratory, more aggressive) [2]. However, if different personalities affiliate, they can share mutual benefits; a concept termed social heterosis [46]. Social heterosis in associations of dissimilar personalities thus can explain why behavioural (and genetic) diversity can evolutionarily persist [46]. Negative frequency-dependent selection is another evolutionary explanation for the existence of behavioural polymorphisms (producers–scroungers, hawks– doves and leaders–followers), and has been shown to maintain diversity in group personality composition [47]. Our findings bring evidence to these potential explanations. Living in social groups with diverse composition can provide benefits in terms of reduced physiological stress. Because physiological state in one life-history stage can have carry-over effects on performance in the subsequent life-history stage (e.g. from wintering to the breeding stage, see e.g. the case of American redstart *Setophaga ruticilla* [48,49]), our results might provide a physiological mechanism that could be responsible for the evolutionary maintenance of behavioural diversity in social groups. These findings, thus, bring helpful insight into the study of social evolution, which is a fundamental question in biology and has implications for human work teams. Although the role of group composition in human team performance is still contentious [10], there is some evidence showing that heterophily is advantageous in project groups that are less stable in time and are engaged in creative tasks, but disadvantageous in production groups that are stable in time and are engaged in routine tasks [12].

### Ethics

All birds participating in the study were released at the site of capture in good health on day 18. The study complies with the ethical guidelines of the BabeLJ-Bolyai University (permit no. 30792) and the European laws regarding animal welfare, and adheres to the ASAB guidelines for the use of animals in behavioural research.

## Data accessibility

All data and analysis code supporting the results are deposited in Dryad Digital Repository (https://doi.org/10.5061/dryad.08kprr51z) [42].

## Authors’ contributions

C.I.V., A.F., and Z.Ba. conceived and designed the study. C.I.V., A.F., P.L.P., O.G., J.P., and Z.Be. performed research. C.I.V., J.P., O.G., and Á.Z.L. contributed new reagents/analytic tools. C.I.V., A.F., and Z.Ba. analysed data. C.I.V., A.F., and Z.Ba. drafted the manuscript with major input from Á.Z.L. and P.L.P. All authors contributed revisions and approved the final version of the manuscript. The authors declare no conflict of interest.

## Supporting information

Additional details for Methods and Results

## Acknowledgements

We thank A. Jakab for providing accommodation for A.F. during the experiment, M. Bán for technical assistance, and V. Bókony and F. Bonier for highly useful comments. The constructive comments raised by two anonymous reviewers considerably improved our manuscript.

