## Additional details for Methods and Results for "Social groups with diverse personalities mitigate physiological stress in a songbird"

*Electronic Supplementary Material – ESM*

DOI: 10.1098/rspb.2020.3092

2. Material and methods

(a) Study protocol

The study is based on a large sample of 240 house sparrows. We caught 40 sparrows during each study replicate (1:1 sex ratio). Sparrows were caught with mist nets at a cattle farm near Bălcaciu village, central Transylvania, Romania (46°11’N, 24°3’E) during six capture sessions (9 November 2014, 5 December 2014, 5 January 2015, 23 January 2015, 10 February 2015, and 28 February 2015). Upon capture (day 0), birds were marked with an aluminium ring, and their sex and body mass (± 0.1 g) was recorded. The birds were transported to the campus of the Babeș-Bolyai University, Cluj-Napoca (46°46’N, 23°33’E) and housed in indoor aviaries for 18 days.

(b) Study timeline

The study timeline was the same for all six replicates. We let the birds to habituate to captivity on the day of capture (day 0) and the next two days (days 1–2). They were housed in four indoor aviary rooms (3 m length × 2 m width × 2.5 m height) in which the birds were distributed randomly in groups of equal sizes (10 birds in each room). The aviary rooms were visually separated from each other. To assess the exploratory behaviour of birds in a novel environment, we first transferred them into individual cages in the morning of day 3 and let them to habituate for two days (days 3–4), then tested for exploration on days 5–7 (see below the details). Day 8 was a resting day. At day 9, we measured the body mass and tarsus length (± 0.01 mm) of the birds, and took the pre-treatment blood sample (150–200 µL; see below the methods). Then the birds were allocated according to an *a priori* defined protocol into one of four social treatment groups of 10 birds each (see below). These four social groups of 10 birds in each of the six study replicates were housed in the same four adjacent aviary rooms as mentioned above. Each social group had an even or quasi-even sex ratio (table S1). The social treatment period lasted nine days until day 18, when we measured again the body mass and took a second blood sample to measure the post-treatment physiological condition. On the same day, we released the birds at the site of capture.

**Table S1.** Sex ratio as shown by the sample sizes per each sex (F – female, M – male) per each experimental group per each study replicate. Each group was formed by 10 birds during each study replicate totalling 40 birds per study replicate and 240 birds for the entire study.

|  | social treatment group | | | | | | | |
| --- | --- | --- | --- | --- | --- | --- | --- | --- |
|  | random | | variable | | low | | high | |
|  | F | M | F | M | F | M | F | M |
| replicate #1 | 4 | 6 | 5 | 5 | 6 | 4 | 5 | 5 |
| replicate #2 | 6 | 4 | 6 | 4 | 4 | 6 | 4 | 6 |
| replicate #3 | 5 | 5 | 3 | 7 | 7 | 3 | 5 | 5 |
| replicate #4 | 5 | 5 | 3 | 7 | 7 | 3 | 5 | 5 |
| replicate #5 | 7 | 3 | 4 | 6 | 5 | 5 | 4 | 6 |
| replicate #6 | 7 | 3 | 4 | 6 | 5 | 5 | 4 | 6 |

(c) Housing and ethical note

Birds were transported within max. 4 h from capture into aviaries. To increase the sparrow’s comfort, aviaries were enriched with several perches and one nest box per bird for resting, hiding and roosting, and a water tank was full-time available for bathing. The artificial photoperiod was identical to the natural day–night cycle throughout. Birds were fed *ad libitum* with a seed mixture consisting of ground corn, barley, millet and sunflower, and this diet was supplemented with one grated boiled egg per aviary room every other day [1,2]. Fresh drinking water was provided on a daily basis. None of the birds died during the study and all of them were released at the site of capture in good health.

(d) Exploratory behaviour

When transferred into individual cages (day 3), birds were randomly ordered from 1 to 40 and split into three clusters (first 13, next 14, and last 13 birds). Their exploration test was performed according to this 1–40 order on days 5–7 (one cluster was tested each day). We recorded exploratory behaviour as a well-established axis of personality following the novel environment test of Dingemanse et al. [3]. Sparrows were deprived of food and water for 1 h before the novel environment test started. Their cage was moved to the test room 10 min before the test run and was covered with a dark curtain, so birds were left to calm down in complete darkness and quietness before the test run. The birds entered the test room from their cage through a sliding door after being startled by knocking the wall of the cage, but without being handled or seeing any person. They were tested alone by spending 10 min in the test room (3 m length × 2 m width × 2 m height) that contained four artificial wooden trees with four branches each and arranged symmetrically within the test room. Exploratory behaviour was video recorded through a one-way window with a hand-held video camera (Panasonic HC-V510) between 09:00 and 16:00 (schedule of the test runs: 09:00, 09:30, 10:00, 10:30, 11:00, 11:30, 12:00, 12:30, 13:00, 13:30, 14:00, 14:30, 15:00, and 15:30) by the same person (A.F.). Exploratory score is the total number of hops (performed either on the trees or on the ground) and flights during the 10-min test. The exploratory behaviour was scored by the same person (Z.Be.).

An additional set of 40 birds that were not involved in the social experiment were assessed thrice for their exploratory behaviour in the same novel environment as the 240 experimental birds in order to verify whether this behavioural trait is consistent in time, a prerequisite of personality traits. The timeline and housing condition for these 40 birds were identical with those 240 birds that were involved in the six study replicates (i.e. they were housed under the same conditions and spent the same number of days before the first test and between the consecutive tests). Consistency of the exploratory behaviour was measured by calculating individual repeatability (i.e. separating variation in exploratory score into a within-individual and an among-individuals component) using a linear mixed-effects model (R package ‘rptR’ [4]) as per Nakagawa and Schielzeth [5]. Exploration score was first log(x+1)-transformed and then Z-transformed (i.e. scaled to mean = 0 and standard deviation = 1; [6]). We first built a full model in which exploration score was the dependent variable with sex (male/female), exploration test repeat (first/second/third), aviary room (from one to four) where the birds were kept between test repeats, and their second-order interaction were entered as potential confounding fixed effects, and individual’s ID, test day (three test days; see above), and the novel environment test order (1–40) nested within test day were entered as random factors. The minimal model was obtained by sequentially dropping all the non-significant fixed predictors from the full model until only significant effects remained. Individuals were significantly consistent in their exploratory behaviour across the three exploration test repeats in both the full model and minimal model (full model: *R* = 0.472, s.e. = 0.097, 95% confidence interval = 0.316–0.691, *p* < 0.001; minimal model: *R* = 0.416, s.e. = 0.100, 95% confidence interval = 0.199–0.589, *p* < 0.001).

(e) Social treatment

The social treatment consisted of creating four groups that differed in personality composition: ‘random’ (random subsample of birds of a given replicate), ‘high variance’ (mixture of birds with either low or high exploration scores), ‘low exploratory’ (only birds with low scores), and ‘high exploratory’ (only birds with high scores). For this, we first ranked the 40 birds of each study replicate according to their exploration score in an increasing order (i.e. rank #1 is the least exploratory bird). The ‘random’ group was set up by forming 10 quartets along this rank order (i.e. first quartet consisting of birds ranking #1–4 and the last quartet of birds ranking #37–40), randomizing the order within each quartet, and then choosing the first bird from each quartet. The ‘high variance’ group was set up by reordering the remaining 30 birds, forming 15 duos along this rank order, randomizing the order of birds within each duo, and choosing the first bird from the first five and the last five duos. The remaining 20 birds were reordered once again and the first 10 birds along this exploration rank order formed the ‘low exploratory’ group, while the last 10 birds formed the ‘high exploratory’ group. The goodness of this protocol was *a priori* assessed by generating 40 random exploration scores with uniform, normal or exponential distribution. The group formation protocol worked for each of the three distribution types as the four groups the protocol created differed both according to mean and to variance of exploration scores.

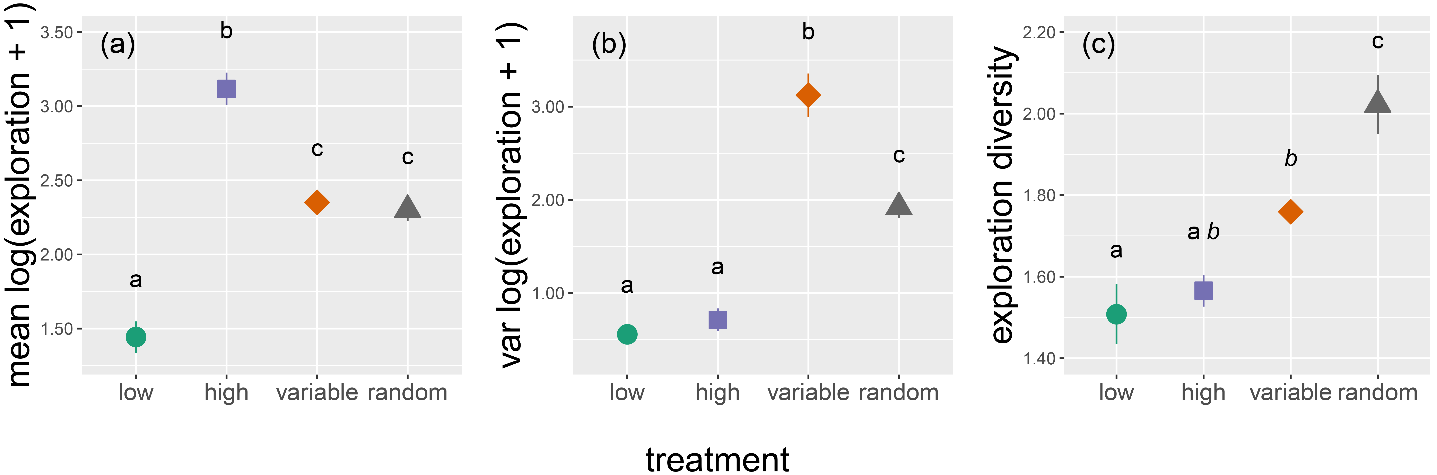

**Figure S1.** Treatment groups differ according to (a) mean, (b) variance and (c) Shannon diversity index of personality (i.e. exploration score). Means ± s.e. are shown on raw data. Different lowercase letters denote significant differences (*p* ≤ 0.050), while similar but italicized letters denote marginal differences (0.050 < *p* < 0.100) between social treatment groups based on post-hoc pairwise comparisons with Tukey-adjusted *p*-values.

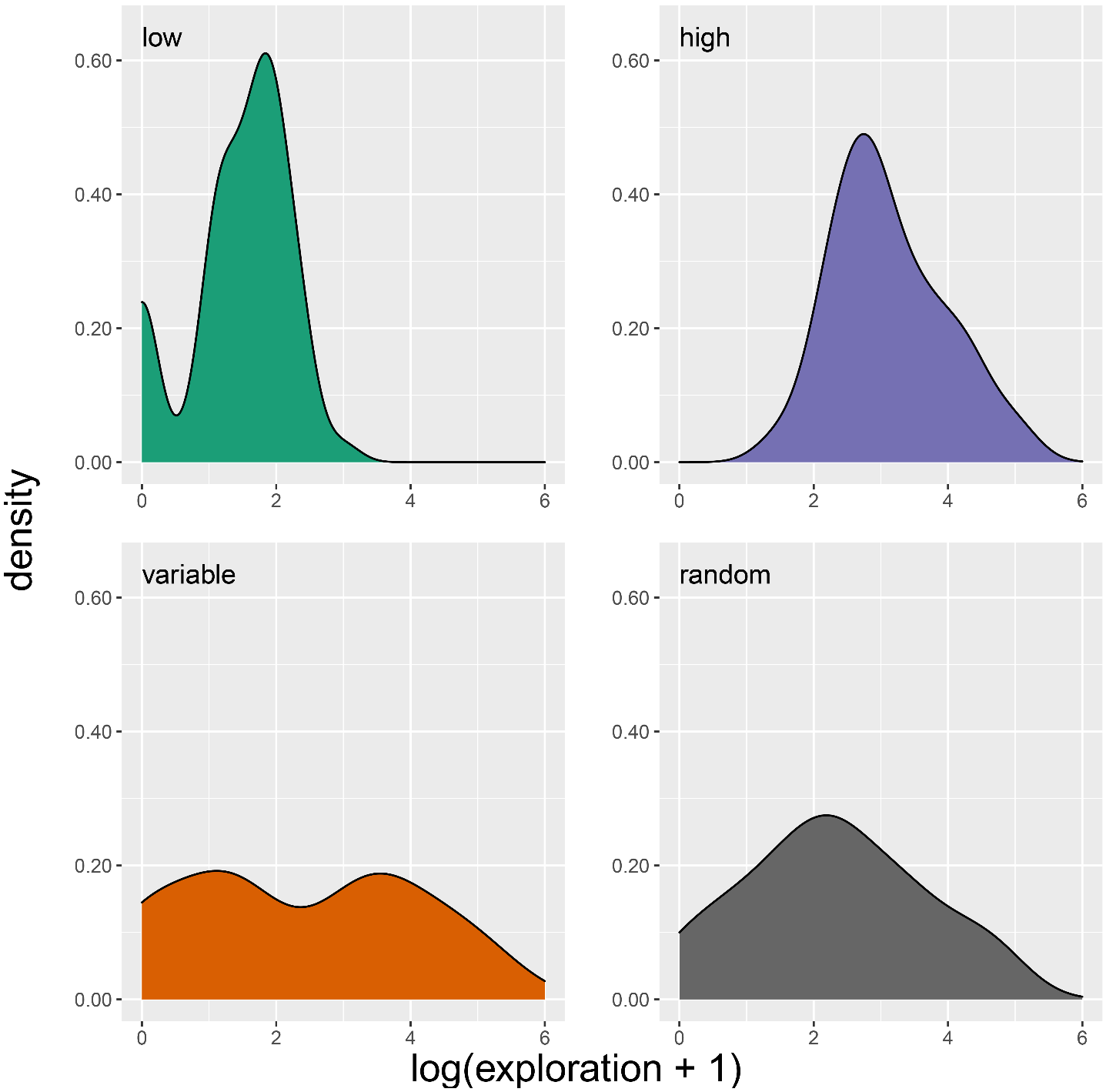

**Figure S2.** Treatment groups differ according to the density distribution of exploration score. The random and variable groups have wider distribution, unimodal in the random group, but bimodal in the variable group. The high and low exploratory groups have narrow distributions at the upper or lower range limit, respectively.

(f) Blood sampling

Blood samples were collected on day 9 and day 18 to assess the physiological state before and after the social treatment period, respectively. Blood samples were collected into heparinized capillaries by puncturing the brachial vein with insulin syringe. A drop of blood was smeared onto a microscope slide for counting leucocytes. The capillaries with blood samples were stored in dark cooling boxes at 4°C for max. 4 h until centrifuged (5 min at 6200 *g*) to separate the plasma and erythrocyte fractions. Plasma was partitioned into aliquots for each physiological parameter and all aliquots were stored at –50°C until the laboratory assay took place.

(g) Physiological parameters

We measured the following five parameters to describe the physiological state of the birds. First, we computed a size-corrected body mass index to characterize the individuals’ body condition (i.e. the relative amount of energy stores in the form of muscle and fat). For this, we used the Scaled Mass Index [7] (for details, see [8]). Second, 50 leukocytes were counted from blood smears by G.O. (for details, see [9,10]). Heterophil-to-lymphocyte ratio was used as an indicator of glucocorticoid-mediated stress response [11]. Because all the leukocytes were heterophils on some smears, heterophil-to-lymphocyte ratio was calculated as heterophils / (heterophils + lymphocytes); thus, a value close to 1 indicates higher physiological stress. Third, oxidative stress was assessed by J.P. and C.I.V. by measuring the amount of oxidative damage to cell membrane phospholipids via the plasma concentration of malondialdehyde, a toxic intermediate of oxidative lipid decomposition (for details, see [12]). Fourth, the level of natural antibodies (agglutination score) and the activity of the complement system (lysis score) as two associated measures of the constitutive innate immune system was assessed by J.P. and C.I.V. via a haemagglutination–haemolysis assay [13] (for details, see [14]). Higher scores mean that the immune system constituents of the plasma can agglutinate or lyse foreign red blood cells at lower concentration (i.e. indicate better immune capacity).

3. Additional results

There was no significant difference among treatment groups in the pre-treatment values of the five physiological variables (body condition, SMI: *χ*^2^ = 0.333, df = 3, *p* = 0.954; heterophil-to-lymphocyte ratio, H/L: *χ*^2^ = 2.441, df = 3, *p* = 0.486; malondialdehyde, MDA: *χ*^2^ = 4.790, df = 3, *p* = 0.188; agglutination: *χ*^2^ = 1.335, df = 3, *p* = 0.721; lysis: *χ*^2^ = 2.007, df = 3, *p* = 0.571).

**Table S2.** Spearman rank correlation coefficients for the pair-wise correlations of the five physiological response variables (SMI – Scaled Mass Index (body condition); H/L ratio – heterophil-to-lymphocyte ratio (indicator of physiological stress); MDA – malondialdehyde (oxidative damage to lipids); agglutination – level of natural antibodies; lysis – activity of the complement system). Upper matrix (i.e. above the diagonal) shows the coefficients for the pre-treatment sampling event, while the lower matrix (i.e. below the diagonal) shows those for the post-treatment sampling event.

|  | SMI | H/L ratio | MDA | agglutination | lysis |
| --- | --- | --- | --- | --- | --- |
| SMI | – | –0.036 | –0.051 | 0.095 | 0.157 |
| H/L ratio | –0.058 | – | 0.038 | –0.037 | –0.146 |
| MDA | 0.014 | –0.057 | – | 0.121 | 0.037 |
| agglutination | 0.038 | –0.17 | 0.23 | – | 0.624 |
| lysis | 0.109 | –0.165 | 0.14 | 0.683 | – |

**Table S3.** Parameter estimates of full models and minimal adequate models of individual responses in physiological state of house sparrows during the social treatment period. Full models contain all the predictors, while minimal models contain the significant predictors and the sampling event × treatment interaction even if not significant (predictor of interest). Statistically significant effects (*t*-value or *z*-value ≥ 2) are marked in bold, while marginally significant effects are marked in italic (1.8 < *t*-value or *z*-value < 2). (a) SMI – Scaled Mass Index (body condition), (b) H/L ratio – heterophil-to-lymphocyte ratio (indicator of physiological stress), (c) MDA – malondialdehyde (oxidative damage to lipids), (e) Agglutination – level of natural antibodies, (f) Lysis – activity of the complement system. Predictors: social treatment (HVG – high variance group, experimental group with high exploratory behaviour variance; HEG – high exploratory group, experimental group of birds with high exploratory behaviour; LEG – low exploratory group, experimental group of birds with low exploratory behaviour; reference level is the random group, experimental group with a random sample of the exploratory behaviour range), S – sex (male is the reference level), SE – sampling event (pre-treatment is the reference level), EB – exploratory behaviour. Random effects: REP – study replication, T – social treatment, ID – individual ID. For random effects, σ^2^ is the residual variance, while τ_00_ is the variance explained by random factors.

(a) SMI

|  | full model | | | min. adequate model | | |
| --- | --- | --- | --- | --- | --- | --- |
| fixed effects | *β* | s.e. | *t*-value | *β* | s.e. | *t*-value |
| intercept | 0.351 | 0.193 | 1.821 | 0.297 | 0.143 | 2.069 |
| **SE** | **0.295** | **0.086** | **3.442** | **0.231** | **0.074** | **3.123** |
| HVG | –0.210 | 0.254 | 0.827 | –0.078 | 0.179 | 0.435 |
| HEG | –0.350 | 0.290 | 1.205 | –0.128 | 0.179 | 0.719 |
| LEG | –0.159 | 0.306 | 0.519 | –0.083 | 0.178 | 0.468 |
| **S** | **–0.524** | **0.254** | **2.063** | **–0.449** | **0.121** | **3.696** |
| EB | 0.082 | 0.149 | 0.552 |  |  |  |
| **SE × HVG** | **–0.276** | **0.105** | **2.624** | **–0.258** | **0.105** | **2.465** |
| **SE × HEG** | **–0.402** | **0.108** | **3.737** | **–0.375** | **0.105** | **3.581** |
| **SE × LEG** | **–0.278** | **0.108** | **2.573** | **–0.292** | **0.105** | **2.789** |
| SE × S | –0.113 | 0.077 | 1.468 |  |  |  |
| SE × EB | 0.022 | 0.042 | 0.530 |  |  |  |
| HVG × S | 0.305 | 0.364 | 0.840 |  |  |  |
| HEG × S | 0.277 | 0.360 | 0.768 |  |  |  |
| LEG × S | 0.162 | 0.364 | 0.445 |  |  |  |
| HVG × EB | –0.022 | 0.164 | 0.132 |  |  |  |
| HEG × EB | 0.009 | 0.237 | 0.039 |  |  |  |
| LEG × EB | –0.099 | 0.256 | 0.389 |  |  |  |
| S × EB | 0.099 | 0.144 | 0.686 |  |  |  |
| random effects | | | | | | |
| σ^2^ | 0.16 | | | 0.16 | | |
| τ_00_ | 0.80 _REP:T:ID_ | | | 0.79 _REP:T:ID_ | | |
|  | 0.00 _REP:T_ | | | 0.00 _REP:T_ | | |
|  | 0.00 _REP_ | | | 0.00 _REP_ | | |
| *n* | 6 REP | | | 6 REP | | |
|  | 4 T | | | 4 T | | |
|  | 240 ID | | | 240 ID | | |
| observations | 480 | | | 480 | | |
| marg. *R*^2^; cond. *R*^2^ | 0.337 / NA | | | 0.282 / NA | | |

(b) H/L ratio

|  | full model | | | min. adequate model | | |
| --- | --- | --- | --- | --- | --- | --- |
| fixed effects | *β* | s.e. | *t*-value | *β* | s.e. | *t*-value |
| intercept | 0.233 | 0.244 | 0.954 | 0.224 | 0.226 | 0.992 |
| **SE** | **–0.567** | **0.193** | **2.943** | **–0.542** | **0.191** | **2.834** |
| HVG | –0.230 | 0.210 | 1.094 | –0.216 | 0.167 | 1.294 |
| HEG | –0.167 | 0.234 | 0.714 | –0.184 | 0.167 | 1.104 |
| LEG | –0.169 | 0.243 | 0.697 | –0.194 | 0.166 | 1.166 |
| S | –0.179 | 0.196 | 0.912 | –0.152 | 0.119 | 1.278 |
| EB | –0.001 | 0.114 | 0.012 |  |  |  |
| SE × HVG | *0.470* | *0.236* | *1.991* | *0.468* | *0.236* | *1.979* |
| **SE × HEG** | **0.559** | **0.241** | **2.316** | **0.617** | **0.236** | **2.616** |
| **SE × LEG** | **0.494** | **0.242** | **2.035** | *0.428* | *0.235* | *1.821* |
| **SE × S** | **0.372** | **0.172** | **2.160** | *0.329* | *0.168* | *1.961* |
| SE × EB | 0.103 | 0.095 | 1.086 |  |  |  |
| HVG × S | 0.012 | 0.255 | 0.046 |  |  |  |
| HEG × S | 0.128 | 0.253 | 0.505 |  |  |  |
| LEG × S | –0.138 | 0.256 | 0.542 |  |  |  |
| HVG × EB | –0.025 | 0.114 | 0.220 |  |  |  |
| HEG × EB | –0.094 | 0.166 | 0.563 |  |  |  |
| LEG × EB | –0.056 | 0.180 | 0.312 |  |  |  |
| S × EB | –0.063 | 0.101 | 0.620 |  |  |  |
| random effects | | | | | | |
| σ^2^ | 0.83 | | | 0.83 | | |
| τ_00_ | 0.01 _REP:T:ID_ | | | 0.00 _REP:T:ID_ | | |
|  | 0.00 _REP:T_ | | | 0.00 _REP:T_ | | |
|  | 0.20 _REP_ | | | 0.20 _REP_ | | |
| *n* | 6 REP | | | 6 REP | | |
|  | 4 T | | | 4 T | | |
|  | 240 ID | | | 240 ID | | |
| observations | 480 | | | 480 | | |
| marg. *R*^2^; cond. *R*^2^ | 0.024 / 0.222 | | | 0.020 / 0.209 | | |

(c) MDA

|  | full model | | | min. adequate model | | |
| --- | --- | --- | --- | --- | --- | --- |
| fixed effects | *β* | s.e. | *t*-value | *β* | s.e. | *t*-value |
| intercept | –0.043 | 0.188 | 0.229 | 0.037 | 0.146 | 0.253 |
| SE | –0.210 | 0.207 | 1.013 | –0.203 | 0.178 | 1.139 |
| HVG | 0.228 | 0.225 | 1.013 | 0.188 | 0.178 | 1.053 |
| HEG | 0.009 | 0.253 | 0.036 | –0.182 | 0.179 | 1.016 |
| LEG | 0.127 | 0.261 | 0.488 | –0.159 | 0.178 | 0.891 |
| S | 0.115 | 0.210 | 0.547 |  |  |  |
| EB | 0.130 | 0.122 | 1.066 |  |  |  |
| SE × HVG | –0.232 | 0.255 | 0.910 | –0.236 | 0.253 | 0.935 |
| *SE × HEG* | *0.500* | *0.262* | *1.909* | *0.467* | *0.254* | *1.837* |
| **SE × LEG** | **0.529** | **0.261** | **2.023** | **0.577** | **0.252** | **2.288** |
| SE × S | 0.018 | 0.187 | 0.094 |  |  |  |
| SE × EB | –0.067 | 0.103 | 0.654 |  |  |  |
| HVG × S | –0.091 | 0.274 | 0.333 |  |  |  |
| HEG × S | –0.068 | 0.274 | 0.247 |  |  |  |
| LEG × S | –0.430 | 0.274 | 1.567 |  |  |  |
| HVG × EB | –0.048 | 0.124 | 0.390 |  |  |  |
| HEG × EB | –0.303 | 0.181 | 1.674 |  |  |  |
| LEG × EB | –0.006 | 0.193 | 0.030 |  |  |  |
| S × EB | –0.134 | 0.109 | 1.224 |  |  |  |
| random effects | | | | | | |
| σ^2^ | 0.95 | | | 0.95 | | |
| τ_00_ | 0.02 _REP:T:ID_ | | | 0.01 _REP:T:ID_ | | |
|  | 0.00 _REP:T_ | | | 0.00 _REP:T_ | | |
|  | 0.03 _REP_ | | | 0.03 _REP_ | | |
| *n* | 6 REP | | | 6 REP | | |
|  | 4 T | | | 4 T | | |
|  | 240 ID | | | 240 ID | | |
| observations | 471 | | | 471 | | |
| marg. *R*^2^; cond. *R*^2^ | 0.044 / NA | | | 0.031 / NA | | |

(d) agglutination

|  | full model | | | min. adequate model | | |
| --- | --- | --- | --- | --- | --- | --- |
| fixed effects | *β* | s.e. | *z*-value | *β* | s.e. | *z*-value |
| intercept | –0.155 | 0.576 | 3.239 | –0.216 | 0.454 | 3.373 |
| **SE** | *2.711* | *0.533* | *1.871* | **3.028** | **0.445** | **2.487** |
| HVG | 2.876 | 0.591 | 1.787 | 1.662 | 0.457 | 1.111 |
| HEG | 1.799 | 0.646 | 0.909 | 1.118 | 0.473 | 0.236 |
| LEG | 2.573 | 0.667 | 1.417 | 1.495 | 0.460 | 0.874 |
| S (F) | 1.778 | 0.542 | 1.061 |  |  |  |
| EB | 1.037 | 0.311 | 0.117 |  |  |  |
| SE × HVG | –0.672 | 0.619 | 0.643 | 0.647 | 0.603 | 0.722 |
| SE × HEG | –0.696 | 0.639 | 0.567 | 0.699 | 0.620 | 0.577 |
| SE × LEG | –0.629 | 0.642 | 0.722 | 0.603 | 0.606 | 0.834 |
| SE × S | 1.226 | 0.448 | 0.454 |  |  |  |
| SE × EB | 1.039 | 0.247 | 0.157 |  |  |  |
| HVG × S | –0.352 | 0.660 | 1.581 |  |  |  |
| HEG × S | –0.510 | 0.660 | 1.021 |  |  |  |
| *LEG × S* | *–0.290* | *0.670* | *1.845* |  |  |  |
| HVG × EB | –0.728 | 0.300 | 1.060 |  |  |  |
| HEG × EB | –0.807 | 0.432 | 0.496 |  |  |  |
| LEG × EB | –0.737 | 0.453 | 0.675 |  |  |  |
| S × EB | 1.091 | 0.259 | 0.336 |  |  |  |
| random effects | | | | | | |
| σ^2^ | 3.29 | | | 3.29 | | |
| τ_00_ | 0.05 _REP:T:ID_ | | | 0.06 _REP:T:ID_ | | |
|  | 0.00 _REP:T_ | | | 0.00 _REP:T_ | | |
|  | 0.44 _REP_ | | | 0.49 _REP_ | | |
| *n* | 6 REP | | | 6 REP | | |
|  | 4 T | | | 4 T | | |
|  | 237 ID | | | 237 ID | | |
| observations | 474 | | | 474 | | |
| marg. *R*^2^; cond. *R*^2^ | 0.074 / NA | | | 0.052 / NA | | |

(e) lysis

|  | full model | | | min. adequate model | | |
| --- | --- | --- | --- | --- | --- | --- |
| fixed effects | *β* | s.e. | *z*-value | *β* | s.e. | *z*-value |
| intercept | –0.160 | 0.633 | 2.896 | –0.132 | 0.531 | 3.816 |
| SE | 1.874 | 0.587 | 1.071 | 2.321 | 0.495 | 1.701 |
| HVG | –0.995 | 0.661 | 0.008 | –0.859 | 0.555 | 0.274 |
| HEG | –0.626 | 0.772 | 0.607 | –0.474 | 0.619 | 1.204 |
| LEG | 1.652 | 0.741 | 0.677 | 1.138 | 0.533 | 0.243 |
| S | –0.630 | 0.633 | 0.729 |  |  |  |
| EB | 1.682 | 0.363 | 1.432 |  |  |  |
| SE × HVG | 1.412 | 0.728 | 0.473 | 1.289 | 0.703 | 0.361 |
| SE × HEG | 1.439 | 0.793 | 0.458 | 1.186 | 0.771 | 0.221 |
| SE × LEG | 1.347 | 0.726 | 0.410 | 1.508 | 0.678 | 0.606 |
| SE × S | 1.495 | 0.560 | 0.717 |  |  |  |
| SE × EB | –0.803 | 0.299 | 0.734 |  |  |  |
| HVG × S | –0.424 | 0.846 | 1.015 |  |  |  |
| HEG × S | –0.877 | 0.846 | 0.155 |  |  |  |
| LEG × S | –0.499 | 0.773 | 0.899 |  |  |  |
| HVG × EB | –0.580 | 0.368 | 1.481 |  |  |  |
| HEG × EB | –0.391 | 0.598 | 1.573 |  |  |  |
| LEG × EB | –0.778 | 0.512 | 0.489 |  |  |  |
| S × EB | –0.582 | 0.340 | 1.591 |  |  |  |
| random effects | | | | | | |
| σ^2^ | 3.29 | | | 3.29 | | |
| τ_00_ | 0.19 _REP:T:ID_ | | | 0.21 _REP:T:ID_ | | |
|  | 0.00 _REP:T_ | | | 0.00 _REP:T_ | | |
|  | 0.65 _REP_ | | | 0.69 _REP_ | | |
| *n* | 6 REP | | | 6 REP | | |
|  | 4 T | | | 4 T | | |
|  | 237 ID | | | 237 ID | | |
| observations | 474 | | | 474 | | |
| marg. *R*^2^; cond. *R*^2^ | 0.179 / NA | | | 0.111 / NA | | |

**Table S4.** Parameter estimates of full models and minimal adequate models of individual responses in physiological state of house sparrows in relation with the Shannon diversity index of the groups during the social treatment period. Full models contain all the predictors, while minimal models contain the significant predictors and the sampling event × Shannon diversity interaction even if not significant (predictor of interest). Statistically significant effects (*t*-value or *z*-value ≥ 2) are marked in bold, while marginally significant effects are marked in italic (1.8 < *t*-value or *z*-value < 2). (a) SMI – Scaled Mass Index (body condition), (b) H/L ratio – heterophil-to-lymphocyte ratio (indicator of physiological stress), (c) MDA – malondialdehyde (oxidative damage to lipids), (e) Agglutination – level of natural antibodies, (f) Lysis – activity of the complement system. Predictors: SE – sampling event (pre-treatment is the reference level), S – sex (male is the reference level), Sh – Shannon diversity index, EB – exploratory behaviour. Random effects: REP – study replication, T – social treatment, ID – individual ID. For random effects, σ^2^ is the residual variance, while τ_00_ is the variance explained by random factors.

(a) SMI

|  | full model | | | min. adequate model | | |
| --- | --- | --- | --- | --- | --- | --- |
| fixed effects | *β* | s.e. | *t*-value | *β* | s.e. | *t*-value |
| intercept | 0.184 | 0.091 | 2.009 | 0.214 | 0.087 | 2.461 |
| SE | 0.047 | 0.053 | 0.895 | 0.000 | 0.037 | 0.000 |
| **S** | **–0.342** | **0.130** | **2.642** | **–0.429** | **0.120** | **3.561** |
| Sh | 0.085 | 0.089 | 0.949 | 0.005 | 0.063 | 0.085 |
| EB | 0.024 | 0.088 | 0.271 |  |  |  |
| SE × S | –0.095 | 0.076 | 1.247 |  |  |  |
| **SE × Sh** | **0.140** | **0.037** | **3.773** | **0.140** | **0.037** | **3.800** |
| SE × EB | –0.001 | 0.038 | 0.024 |  |  |  |
| S × Sh | –0.153 | 0.121 | 1.271 |  |  |  |
| S × EB | 0.112 | 0.124 | 0.904 |  |  |  |
| random effects | | | | | | |
| σ^2^ | 0.16 | | | 0.16 | | |
| τ_00_ | 0.78 _REP:T:ID_ | | | 0.79 _REP:T:ID_ | | |
|  | 0.00 _REP:T_ | | | 0.00 _REP:T_ | | |
|  | 0.00 _REP_ | | | 0.00 _REP_ | | |
| *n* | 6 REP | | | 6 REP | | |
|  | 4 T | | | 4 T | | |
|  | 240 ID | | | 240 ID | | |
| observations | 480 | | | 480 | | |
| marg. *R*^2^; cond. *R*^2^ | 0.307 / NA | | | 0.259 / NA | | |

(b) H/L ratio

|  | full model | | | min. adequate model | | |
| --- | --- | --- | --- | --- | --- | --- |
| fixed effects | *β* | s.e. | *t*-value | *β* | s.e. | *t*-value |
| intercept | 0.079 | 0.201 | 0.392 | 0.000 | 0.191 | 0.000 |
| SE | –0.170 | 0.120 | 1.415 | 0.000 | 0.084 | 0.000 |
| S | –0.159 | 0.122 | 1.305 |  |  |  |
| Sh | 0.012 | 0.076 | 0.160 | 0.032 | 0.061 | 0.517 |
| EB | –0.042 | 0.074 | 0.566 |  |  |  |
| *SE × S* | *0.340* | *0.173* | *1.971* |  |  |  |
| SE × Sh | –0.069 | 0.084 | 0.817 | –0.066 | 0.084 | 0.787 |
| SE × EB | 0.113 | 0.087 | 1.303 |  |  |  |
| S × Sh | 0.039 | 0.085 | 0.460 |  |  |  |
| S × EB | –0.010 | 0.087 | 0.115 |  |  |  |
| random effects | | | | | | |
| σ^2^ | 0.84 | | | 0.84 | | |
| τ_00_ | 0.00 _REP:T:ID_ | | | 0.00 _REP:T:ID_ | | |
|  | 0.00 _REP:T_ | | | 0.00 _REP:T_ | | |
|  | 0.20 _REP_ | | | 0.20 _REP_ | | |
| *n* | 6 REP | | | 6 REP | | |
|  | 4 T | | | 4 T | | |
|  | 240 ID | | | 240 ID | | |
| observations | 480 | | | 480 | | |
| marg. *R*^2^; cond. *R*^2^ | 0.011 / NA | | | 0.001 / NA | | |

(c) MDA

|  | full model | | | min. adequate model | | |
| --- | --- | --- | --- | --- | --- | --- |
| fixed effects | *β* | s.e. | *t*-value | *β* | s.e. | *t*-value |
| intercept | 0.019 | 0.113 | 0.172 | –0.001 | 0.092 | 0.007 |
| SE | –0.022 | 0.130 | 0.173 | –0.001 | 0.090 | 0.015 |
| S | –0.050 | 0.131 | 0.379 |  |  |  |
| Sh | 0.032 | 0.081 | 0.389 | 0.054 | 0.065 | 0.832 |
| EB | 0.015 | 0.081 | 0.192 |  |  |  |
| SE × S | 0.042 | 0.188 | 0.221 |  |  |  |
| **SE × Sh** | **–0.217** | **0.091** | **2.371** | **–0.224** | **0.091** | **2.470** |
| SE × EB | –0.065 | 0.095 | 0.691 |  |  |  |
| S × Sh | 0.039 | 0.092 | 0.426 |  |  |  |
| S × EB | –0.044 | 0.095 | 0.460 |  |  |  |
| random effects | | | | | | |
| σ^2^ | 0.97 | | | 0.96 | | |
| τ_00_ | 0.00 _REP:T:ID_ | | | 0.00 _REP:T:ID_ | | |
|  | 0.00 _REP:T_ | | | 0.00 _REP:T_ | | |
|  | 0.03 _REP_ | | | 0.03 _REP_ | | |
| *n* | 6 REP | | | 6 REP | | |
|  | 4 T | | | 4 T | | |
|  | 240 ID | | | 240 ID | | |
| observations | 471 | | | 471 | | |
| marg. *R*^2^; cond. *R*^2^ | 0.019 / 0.044 | | | 0.016 / NA | | |

(d) agglutination

|  | full model | | | min. adequate model | | |
| --- | --- | --- | --- | --- | --- | --- |
| fixed effects | *β* | s.e. | *z*-value | *β* | s.e. | *z*-value |
| intercept | –0.314 | 0.364 | 3.189 | –0.277 | 0.335 | 3.828 |
| **SE** | **1.957** | **0.305** | **2.199** | **2.208** | **0.219** | **3.615** |
| S | –0.808 | 0.330 | 0.645 |  |  |  |
| Sh | –0.767 | 0.217 | 1.222 | –0.805 | 0.181 | 1.194 |
| EB | –0.797 | 0.197 | 1.151 |  |  |  |
| SE × S | 1.284 | 0.438 | 0.572 |  |  |  |
| SE × Sh | 1.306 | 0.227 | 1.177 | 1.318 | 0.226 | 1.220 |
| SE × EB | 1.051 | 0.222 | 0.222 |  |  |  |
| S × Sh | 1.134 | 0.226 | 0.555 |  |  |  |
| S × EB | 1.190 | 0.225 | 0.773 |  |  |  |
| random effects | | | | | | |
| σ^2^ | 3.29 | | | 3.29 | | |
| τ_00_ | 0.06 _REP:T:ID_ | | | 0.07 _REP:T:ID_ | | |
|  | 0.00 _REP:T_ | | | 0.00 _REP:T_ | | |
|  | 0.45 _REP_ | | | 0.48 _REP_ | | |
| *n* | 6 REP | | | 6 REP | | |
|  | 4 T | | | 4 T | | |
|  | 237 ID | | | 237 ID | | |
| observations | 474 | | | 474 | | |
| marg. *R*^2^; cond. *R*^2^ | 0.060 / NA | | | 0.052 / NA | | |

(e) lysis

|  | full model | | | min. adequate model | | |
| --- | --- | --- | --- | --- | --- | --- |
| fixed effects | *β* | s.e. | *z*-value | *β* | s.e. | *z*-value |
| intercept | –0.157 | 0.438 | 4.235 | –0.113 | 0.414 | 5.267 |
| **SE** | **2.383** | **0.344** | **2.523** | **2.851** | **0.260** | **4.022** |
| S | –0.455 | 0.440 | 1.789 |  |  |  |
| Sh | 1.067 | 0.250 | 0.258 | –0.927 | 0.225 | 0.335 |
| EB | –0.938 | 0.231 | 0.279 |  |  |  |
| SE × S | 1.436 | 0.542 | 0.668 |  |  |  |
| SE × Sh | 1.124 | 0.270 | 0.434 | 1.087 | 0.268 | 0.311 |
| SE × EB | –0.778 | 0.270 | 0.929 |  |  |  |
| S × Sh | –0.737 | 0.272 | 1.124 |  |  |  |
| S × EB | –0.777 | 0.278 | 0.907 |  |  |  |
| random effects | | | | | | |
| σ^2^ | 3.29 | | | 3.29 | | |
| τ_00_ | 0.16 _REP:T:ID_ | | | 0.23 _REP:T:ID_ | | |
|  | 0.00 _REP:T_ | | | 0.00 _REP:T_ | | |
|  | 0.63 _REP_ | | | 0.65 _REP_ | | |
| *n* | 6 REP | | | 6 REP | | |
|  | 4 T | | | 4 T | | |
|  | 237 ID | | | 237 ID | | |
| observations | 474 | | | 474 | | |
| marg. *R*^2^; cond. *R*^2^ | 0.128 / NA | | | 0.078 / NA | | |

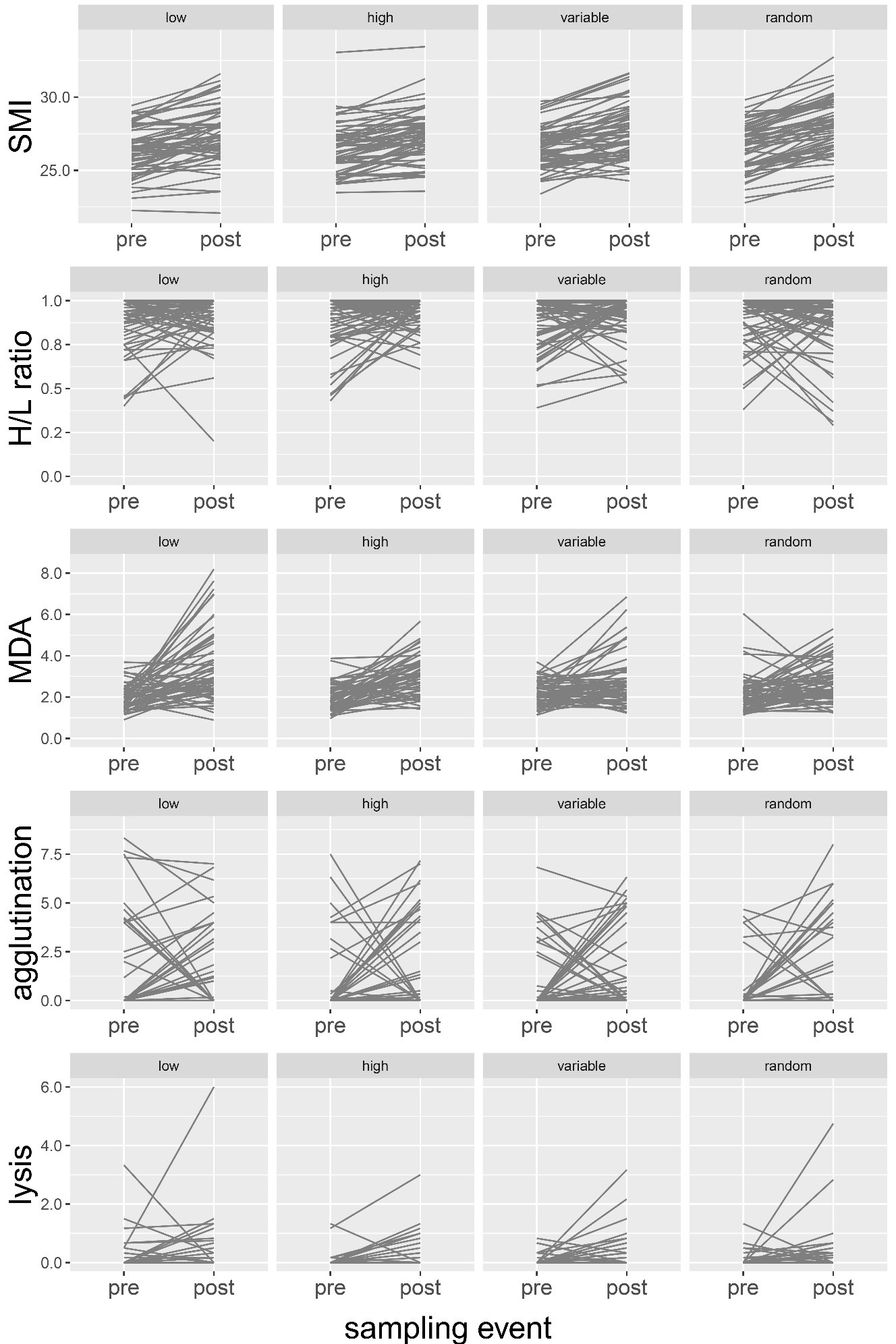

**Figure S3.** Individual reaction norms of physiological responses to social treatment. Each line denotes an individual (*N* = 60 per group) connecting the pre-treatment value (sampling event = pre) with the post-treatment value (sampling event = post). Treatment groups: low = low exploratory group, high = high exploratory group, variable = variable group, random = random group.
